# Identification and characterization of *Botrylloides* (Styelidae) species from Aotearoa New Zealand coasts

**DOI:** 10.1101/2021.09.08.459371

**Authors:** Berivan Temiz, Rebecca M. Clarke, Mike Page, Miles Lamare, Megan J. Wilson

## Abstract

Ascidians are marine filter feeder primitive chordates. *Botrylloides* ascidians possess diverse biological abilities like whole-body regeneration (WBR), hibernation/aestivation, blastogenesis, metamorphosis, and natural chimerism. However, the absence of distinctive morphological features often makes identification problematic. *Botrylloides diegensis* is an ascidian that has been misidentified in previous studies and is recorded in GenBank as *Botrylloides leachii* due to the high morphological similarity between the sister species. More available sequences and strategies around identification would help resolve some of the confusion currently surrounding its ambiguous nature. We collected several *Botrylloides* samples from 7 locations around New Zealand - Dunedin, Christchurch, Picton, Nelson, Whangateau, New Plymouth and Invercargill - and barcoded the species based on Cytochrome Oxidase I, Histone 3, 18S, and 28S ribosomal RNA markers. Network and Bayesian trees confirmed the presence of three *Botrylloides* species: *B. diegensis, B. jacksonianum*, and *B. aff. anceps*. Additionally, morphotypes of three species were investigated regarding zooid size, area, tentacle numbers and colonial arrangement.

## INTRODUCTION

Ascidiacea includes approximately 3000 identified species that are filter-feeding marine invertebrates, many with global distribution (Shenkar & Swalla, 2011). For example, *Botryllus* (Gaertner, 1774) and *Botrylloides* (Milne-Edwards, 1841) are sessile colonial styelid tunicates found within the intertidal and shallow subtidal zones (Kocot et al., 2018). The colony is comprised of numerous genetically identical zooids inside the tunic, an extracellular gelatinous matrix. The tunic contains a cellulose-like substance called tunicine and covers the colonial systems, and protects from the outer environment as a barrier (Belton et al., 1989).

Compound tunicates belong to the sedentary group, which possesses two life stages. The first is a free-swimming tadpole larva without any extrinsic feeding ability until it attaches to a substratum. These free-living larvae can attach to many substrates, such as stones, ship hulls, mussels, and seaweed. The sedentary life stage starts after the attachment. It is followed by metamorphosis resulting in the loss of chordate features such as the notochord, external pharyngeal gill slits, nerve cord, and post-anal tail (Blanchoud, Rinkevich, et al., 2018). After metamorphosis, the individual adults (zooids) can reproduce asexually through a weekly asexual budding cycle called blastogenesis (Berrill, 1947). During this cycle, individuals form buds from their endothelial epithelium, generating new zooids while the original zooid is absorbed.

Tunicates are more closely related to the subphylum Vertebrata than Cephalochordata and, therefore, are of interest in evolution studies (Delsuc et al., 2006). *Botrylloides* ascidians additionally have several notable abilities. Firstly, they can undergo whole-body regeneration (WBR), forming a new adult from a vascular fragment following the loss of all zooids (Rinkevich et al., 1995). Secondly, colonies can enter a dormancy period in suboptimal environmental conditions, where all the zooids are absorbed, and only a compact vasculature is left (Burighel et al., 1976). Thirdly, natural chimerism, the fusion of two contacting colonies that share similar histocompatibility alleles, also known as allorecognition, results from recognizing the self from xeno-recognition (Manni et al., 2019; Rinkevich, 2005; Rinkevich et al., 2007). WBR, blastogenesis, hibernation/aestivation, and natural chimerism are reported only within the Styelidae ascidians. To date, examples of WBR have been described for *Botrylloides violaceus* (Oka, 1927)*, Botrylloides diegensis* (Ritter & Forsyth, 1917), *Botryllus primigenus* (Oka, 1928)*, Botrylloides anceps* (Herdman, 1891) and *Botryllus schlosseri*, Pallas (1766) (Blanchoud, Rinkevich, et al., 2018; Karahan et al., 2022; Rosner et al., 2019). *B. diegensis* can regenerate continuously from a little vascular tissue that includes at least ~200 cells in as little as 10 days (Blanchoud, Rutherford, et al., 2018; Rinkevich et al., 1995). Investigating these biological differences between the tunicate species gives highly profound information about their evolutionary history.

Ascidians have been studied ecologically and regularly monitored to determine if they are either indigenous or invasive; however, due to the absence of solid biogeographical or historical evidence, it is challenging to develop a list of their origin or introductions. Nevertheless, this is important as spatial competition is a vital phenomenon for sessile species’ survival, which can harm native species (Zhan et al., 2015). Due to the increase in sea transportation and the construction of shipping channels, new introductions have increased in the last century, altering habitat structures and biodiversity (Carman et al., 2010). For example, *B. schlosseri* and *B. violaceus* are known to be invasive and reported to affect aquaculture negatively (Karahan et al., 2016). Once these species are introduced to a new habitat, they can overgrow and dominate the available spaces, including the mobile areas, such as the outer layer of crustaceans, which might adversely affect the animal’s mobility. *B. diegensis* is commonly seen in the intertidal zone throughout New Zealand’s coasts and is thought to have originated from the Western or Southern Pacific, while *B. leachii* is stated to have Mediterranean origins (Carlton, 2009; Page & Kelly, 2013; Page et al., 2014; Viard et al., 2019). *B. diegensis* is thought to be introduced to the Atlantic and Northern Pacific (Carlton, 2009; Viard et al., 2019), unlike *B. leachii*, which was stated to be non-indigenous in Australia and Tasmania (Shenkar & Swalla, 2011).

Ascidians contain crypticity due to morphological plasticity and slight differences in anatomical distinctions between species. The identification of *Botrylloides* species has been particularly challenging. Their separation from sister species is unclear due to a lack of morphological divergence or a defined distance for inter/intra-species delineation (Reem et al., 2018; Viard et al., 2019). The sister species which lack this are *Botrylloides leachii, Botrylloides violaceus, Botrylloides niger* (Herdman, 1886), or *Botrylloides diegensis* (Brunetti, 2009; Reem et al., 2018; Salonna et al., 2021; Viard et al., 2019). *Botrylloides perspicuus* (Herdman, 1886)*, Botrylloides giganteus* (Pérès, 1949), and *Botrylloides pizoni* (Brunetti & Mastrototaro, 2012) have similar colonial and zooidal features, although they are distinct species (Rocha et al., 2019). The similarities between these sister *Botrylloides* species have resulted in ambiguous identification and sometimes misidentification, demonstrating the necessity for clear taxonomical identification.

DNA barcoding is a powerful tool and, when combined with morphological data, enables the identification of species, including the genus *Botrylloides* (Miralles et al., 2016; Reem et al., 2018; Rocha et al., 2019). The selection of molecular markers to estimate the divergence is essential. These studies often use mitochondrial cytochrome oxidase subunit I (COI), a polymorphic but conserved region, and other nuclear gene markers such as 12S rRNA, 16S rRNA, 18S rRNA, and 28S rRNA genes can also be used to increase the resolution of identification (Reem et al., 2018). Improving the quality and quantity of database sequences will play an essential role in future barcoding approaches and improving global biodiversity monitoring. For example, barcoding combined with the morphological investigation has found that *Ciona robusta* was erroneously assigned as *Ciona intestinalis*, a different species (Brunetti et al., 2015). In other cases, mitogenomics, a whole mitochondrial barcoding technique, has been used to separate species. For example, a recent study found that a European clade of *Botryllus schlosseri* was a new species called *Botryllus gaiae* (Brunetti et al., 2020). Significantly, a recent study suggested all the GenBank sequences of *Botrylloides leachii* were incorrectly assigned, and these sequences belong to *Botrylloides diegensis* (Viard et al., 2019). Furthermore, it was stated that the spoked wheel morphology in the vicinity of the zooidal buccal siphon is present in *B. leachii* but absent in *B. diegensis*. Considering all these three species are commonly used model invertebrate chordates due to their prominent phylogenetic position makes accurate identification essential.

Our group studies regeneration in *Botrylloides*; we had assigned this species as *B. leachii* based on previous publications and comparisons to GenBank sequences. However, considering the recent studies, we aimed to use DNA barcoding combined with morphology to identify *Botrylloides* species from Aotearoa New Zealand coasts.

## MATERIALS & METHODS

### Sampling Area

In total 40 samples were collected from 7 different intertidal zones around New Zealand (Invercargill, Dunedin, Christchurch, Nelson, Picton, Whangateau, New Plymouth). Samples were preserved in absolute ethanol, and the samplings were performed from July 2020 to February 2021. Sampling details, including the collection date, haplotype information, sampling stations, their coordinates and the sampling dates, are listed for all collected samples (Table S1). Samples were taken from the first few meters of the ocean surface, except for a couple of samples acquired from different depths via diving. The collection was based on morphological identification at the sampling sites based on the general knowledge of *Botrylloides* ascidian morphology. A colony fragment is taken using a single-edged razor blade. For animal breeding, living tunicate tissue fragments are attached to 5×7.5 cm glass slides (Zondag et al., 2016). The slides are placed in tanks filled with filtered salt water that is constantly aerated. Animals are fed regularly with a shellfish diet, and their water is replaced every two days.

### Morphological Examination

Collected *Botrylloides* ascidians were photographed and examined morphologically for the zooid arrangement and colour. Some samples were only taken for molecular analysis; therefore, not all the colonies’ images were taken. Orange and brown-white morphotypes of *B. diegensis*, purple morphotypes of *B. jacksonianum* and purple *B. aff.anceps* were investigated for their tentacle numbers, size and area of the zooids. Size and area of zooids were measured using cellSens Standard 1.5 imaging suite using Nikon SMZ745 dissection microscope. Only some of the living morphotypes of the three species were checked for their zooid sizes and areas.

### Molecular Analysis

DNA extraction was performed based on Gemmell and Akiyama (1996). The alcohol was removed from the samples, and 300 μl lysis buffer (100 mM NaCl, 50 mM TrisCl, 1% SDS, 50 mM EDTA pH 8) was added to each tube. Proteinase K (20 mg/ml) was added to a final concentration of 100 μg/ml. The colonies were homogenized and left to incubate for 2 h at 50°C. Following tissue digestion, 300 μl of 5 M LiCl was added to the tubes. The lysate was mixed for 1 min through inversion. Next, 600 μl of chloroform was added, and the samples were left on a rotating wheel for 30 min. The samples were spun for 15 min at max speed. The supernatant was placed in a new tube. Two volumes of absolute ethanol were added to the tubes and were inverted several times. DNA was precipitated by centrifuging at max speed for 30 min. The supernatant was discarded, and the pellet was washed with 70% ethanol and centrifuged at max speed for 5 min. Excess ethanol was removed, and the pellets were left to air dry for 10 min. Finally, 100-200 μl TE buffer (10 mM TrisCl, 1 mM EDTA – pH 7.5) was used to resuspend the pellet, and the tubes were left overnight at 4°C. DNA samples were stored at −20°C.

DreamTaq Green PCR Master Mix (ThermoFisher) was used with 10 ng/μl diluted genomic DNA. PCR was performed as following: 1) 95°C – 3min; 1 cycle 2) 95°C - 30 sec, 55°C – 30 sec, 72°C - 1 min for 35 cycles 3) 72°C - 10min. The primers used in this study are given in Table 1. The PCR products are cleaned with ExoSAP-IT™ PCR Product Cleanup Reagent (ThermoFisher). Five μl of post-PCR product is mixed with 2 μl of ExoSAP reagent and incubated at 37°C for 15 min and then 80°C for 15 min.

**Table 1.**
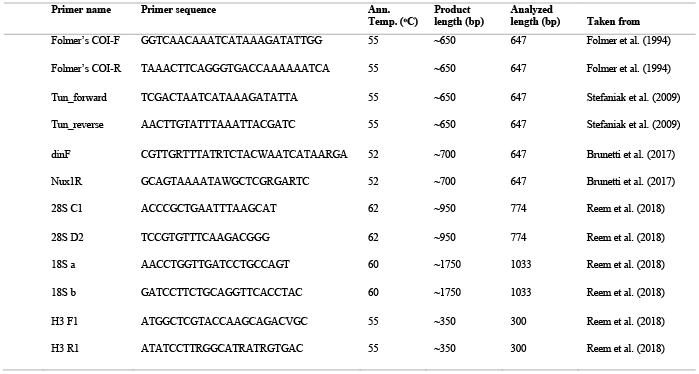
Primer sequences, annealing temperature, expected product length for COI, H3, 18S, and 28S barcodes used in this study.

Two samples (*B. aff. anceps* and *B. jacksonianum*) required cloning the PCR into a vector for sequencing. First, the DNA fragments were amplified using primers from gDNA. Then, the fragments were ligated into pCRII-TOPO (Life Technologies). Next, competent *E. coli* cells were transformed with the plasmid and selected via blue-white screening. Finally, the plasmids were isolated and sequenced. Otago University Sequencing Facility sequenced all samples via sanger sequencing. The tubes were prepared as 3.2 pmol M13 forward primer and 4 μl of cleaned post-PCR template per sample.

All sequences were checked based on their chromatogram data and trimmed if required so that only the confident base calls were used for the alignment. The current study sequences were uploaded to GenBank (Table S2). For comparative analysis, *Botrylloides* COI sequences from GenBank were also added to the Network and Phylogenetic analyses (Table S3). The outgroup species of the COI, H3, 18S and 28S phylogenies are *Symplegma brakenhielmi* (Accession no: LS992554), *Styela clava* (Accession no: XR_005568549), *Symplegma viride* (Accession no: DQ346655) and *Ciona intestinalis* (Accession no: AF212177) respectively.

Forward and reverse sequences are aligned, edited, and trimmed using BioEdit version 7 per marker for Multiple alignments (Hall, 1999) and Clustal X (Thompson et al., 1997). Finally, COI haplotypes are identified using NETWORK version 10 with the median-joining algorithm (Bandelt et al., 1999).

Bayesian probabilities and branch lengths were calculated for all loci via MrBayes 3.2 (Ronquist et al., 2012). We used the HKY+I+G model for COI, HKY+G for H3, K80+G for 18S and GTR+G model for 28S phylogenies (Hasegawa et al., 1985; Kimura, 1980; Tavaré, 1986). The best model selection calculations were computed via jmodeltest2 with the AICs and BIC methods for the Bayesian phylogenies (Darriba et al., 2012). The posterior probabilities of mitochondrial and nuclear sequences were analyzed with two independent runs of Monte-Carlo Markov. At least a million generations were calculated with a sampling frequency of 1000 for each generation for nuclear genes while over 15 million generations were generated for the COI tree. The split frequencies of all the phylogeny calculations were lower than 0.01. The Bayesian trees were optimized using FigTree 1.4.4 (Rambaut, 2018).

Operational taxonomic units (OTUs) were selected based on the analyses through the Automatic Barcode Gap Discovery (ABGD) method of Assemble Species by Automatic Partitioning (ASAP) (Puillandre et al., 2021). Jukes-Cantor and Kimura 2 parameter-based estimations were calculated with ASAP (Jukes & Cantor, 1969; Kimura, 1980).

## RESULTS

### Morphological observations

*Botrylloides* zooids are lined side by side as branching double-row systems, also called *“leachii”* type, with their dorsal lamina facing the surrounding environment. Different *B. diegensis* morphs from New Zealand coasts were observed, including orange, brown-orange, brown, and brown-white or purple-white (Fig. 1) (Table S4). It is difficult to differentiate the brown-white morph from purple-white in some cases; thus, they might be the same, different or can be a transition morph. The zooid sizes and the area of the zooids of *B. diegensis* and *B. affinis anceps* are very similar (~130 um length & ~7500 um^2^ area) (Fig. 2A, B, D, E, F). In comparison, zooids of *Botrylloides jacksonianum* (Herdman, 1899) are larger in size and area (~160 um & ~12800 um^2^) (Fig. 2C, E, F). The shapes of the zooids are similar within *B. diegensis* and *B. affinis anceps*, elliptical egg-like individuals (Fig. 2A, B, D), while *B. jacksonianum* zooids are more nodal and finger-like towards the atrial tongue (Fig. 2C). All zooids are equal in tentacle numbers (4 large, 4 smaller, and 8 petite) (Fig. 2A-D). There are white-pigmented cells on the tentacles of *B. jacksonianum* (Fig. 2C). The two largest-lateral tentacles are distinct at the buccal siphon of *B. affinis anceps* (Fig. 2D).

**Figure 1.**
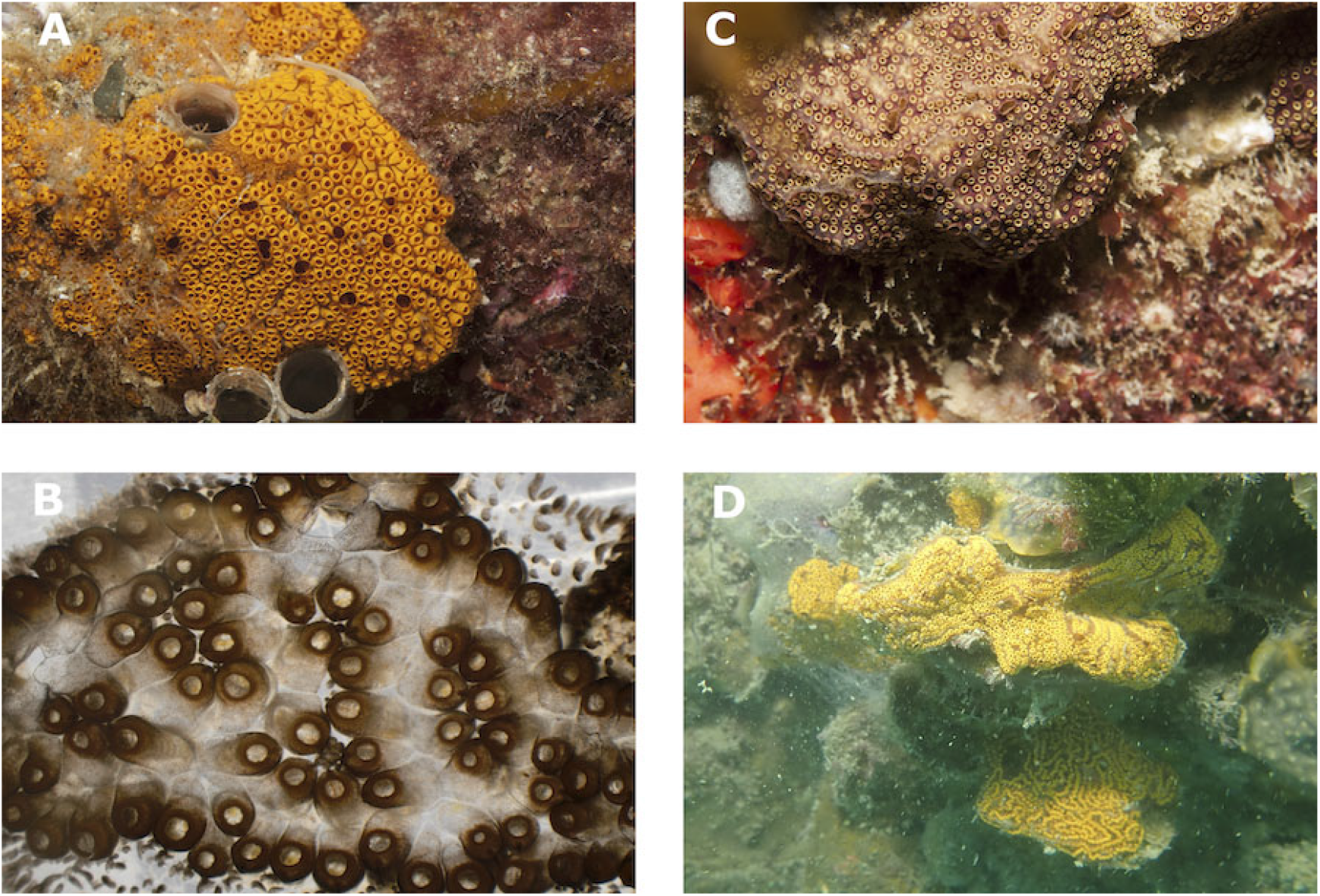
Different color morphs of *Botrylloides diegensis* from Aotearoa New Zealand. **A)** Common orange morph. **B)** A brown-orange *B. diegensis* colony. **C)** Brown-white morph. **D)** Brown-orange colony.

**Figure 2.**
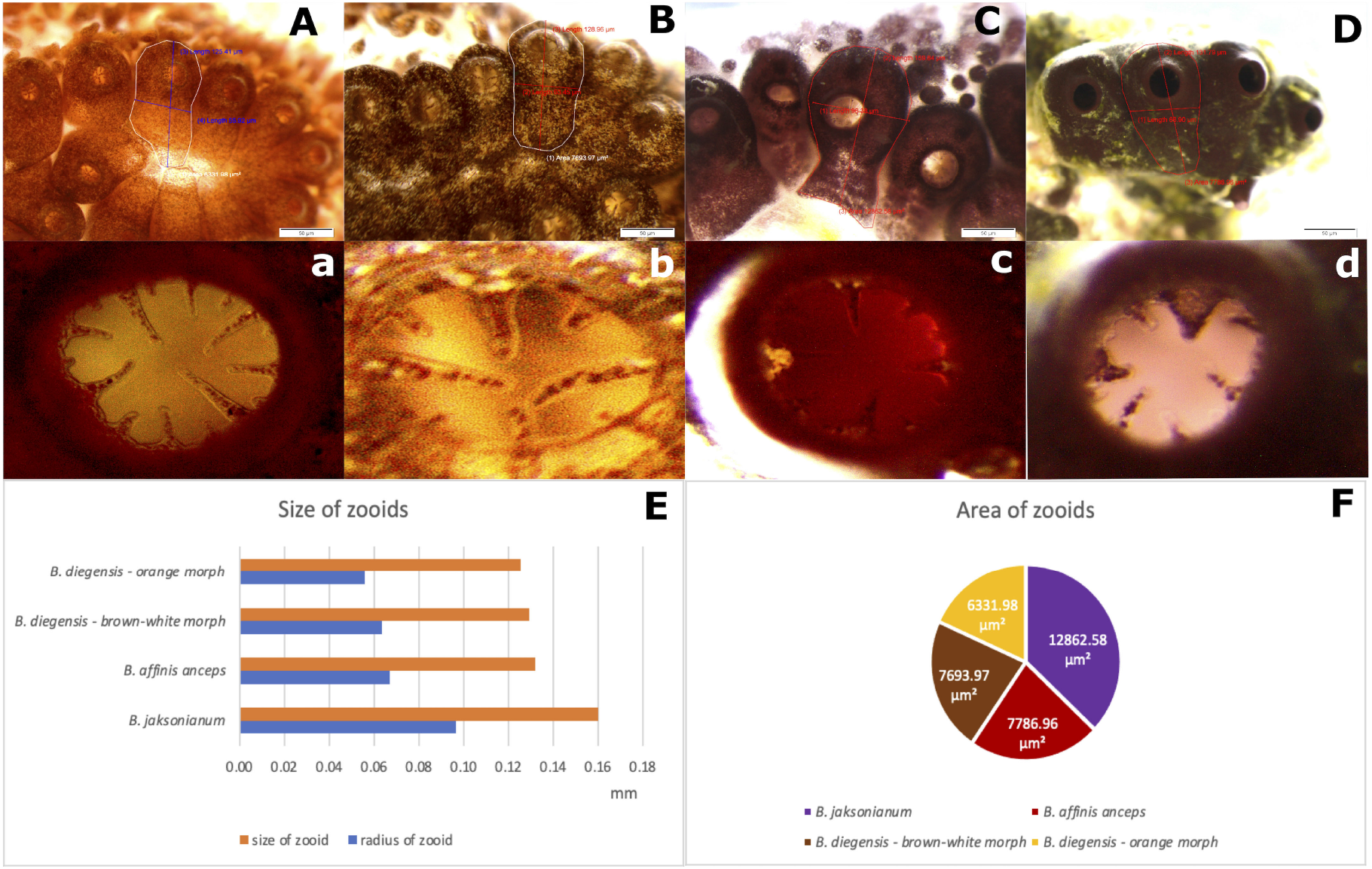
Differences in zooidal size and morphologies of Botryllid species. **A)** Zooid of the orange morph of *B. diegensis*. **a)** Tentacles of the orange morph of *B. diegensis* **B)** Zooid of the brown-white *B. diegensis* colony. **b)** Tentacles of brown-white *B. diegensis*. **C)** Zooid of *B. jacksonianum* **c)** Tentacles of *B. jacksonianum*. **D)** Zooid of *B. affinis anceps* **d)** Tentacles of *B. affinis anceps*. **E)** Bar chart summarising the size and radius of different botryllid zooids. **F)** Pie chart represents the area of the given zooids. Example measurements are shown in the earlier panels A-D. Scales are equal to 50 μm.

### Phylogeny and haplotype diversity

A phylogeny based on COI locus was constructed to evaluate their taxonomy (Fig. 3). Whangateau and Invercargill colonies clustered with *B. cf. anceps* COI barcode sequence from Australia (Accession no: MT873573) and the sample from Dunedin grouped within *B. jacksonianum* from Australia (Accession no: MT873572). While the distance is high within the *B. anceps* cluster and thus identified as *B. affinis anceps*, the colony from Dunedin is nominated as *B. jacksonianum* due to the low mismatch ratio (<1%). The rest of the sequences clustered with *B. diegensis* sequences from GenBank (Fig. 3).

**Figure 3.**
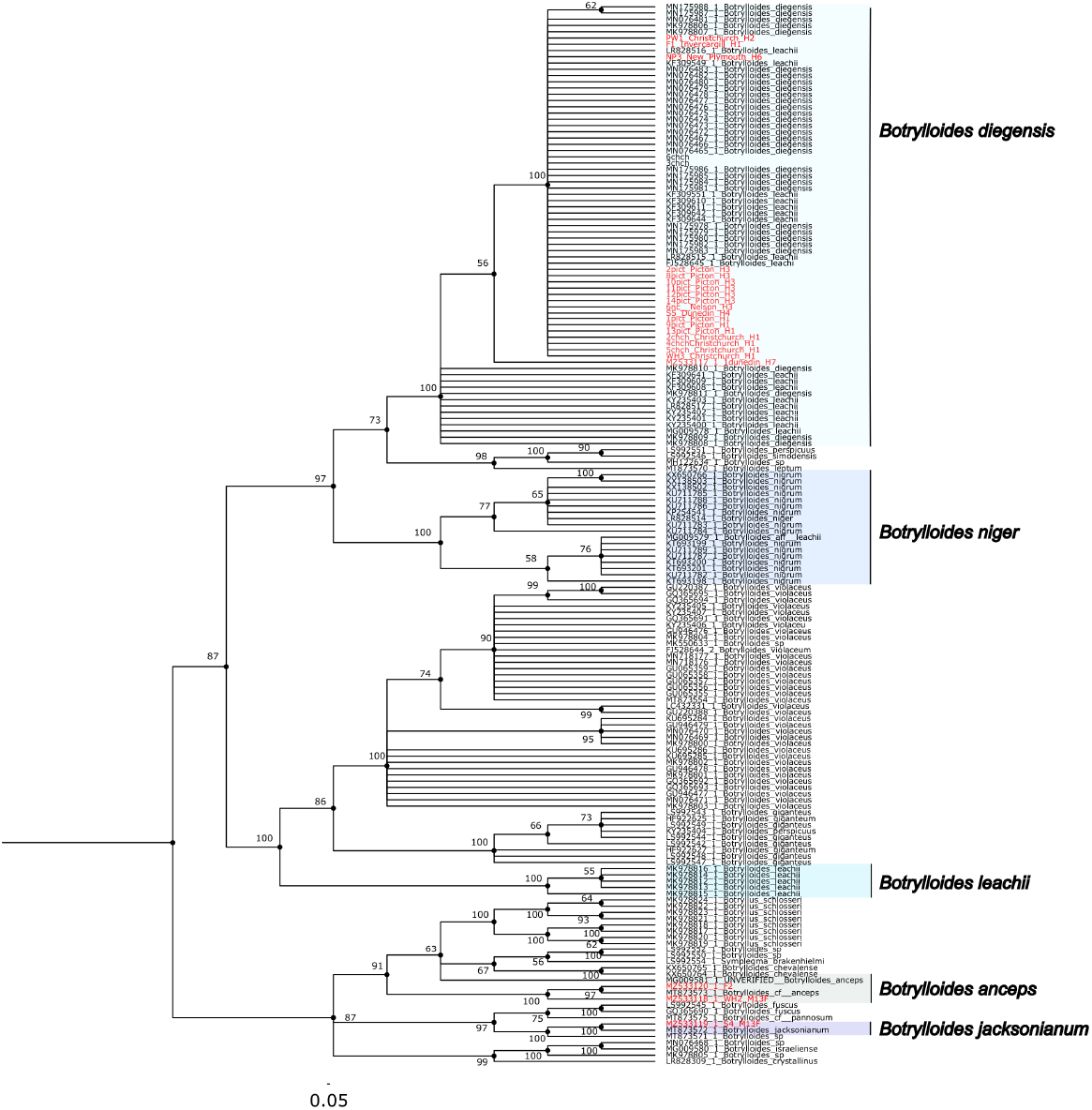
Bayesian tree of *Botrylloides* based on mitochondrial COI locus. Current study sequences and all database *Botrylloides* sequences from GenBank were used to construct the tree. Two independent runs were executed with Monte-Carlo Markov Chains. Ten million generations were measured, with a sampling frequency of 1000 for each generation. Split frequencies were lower than 0.01. Red sequences indicate the current study sequences.

Using ASAP analysis to partition species, 23 operational taxonomic units (OTUs) were identified (Figure 4; 153 sequences). All *B. diegensis* samples of this study clustered with the database *B. diegensis* samples in the same OUT, along with some of the database sequences previously misidentified identified as *B. leachii* (71 sequences in total). Confirmed *B. leachii* sequences formed a separate OTU (Fig. 4). *B. jacksonianum* was another OTU. *B. aff. anceps* samples of this study were two distinct OTUs closely aligned with the *B. cf. anceps* database OTU (MT873573).

**Figure 4.**
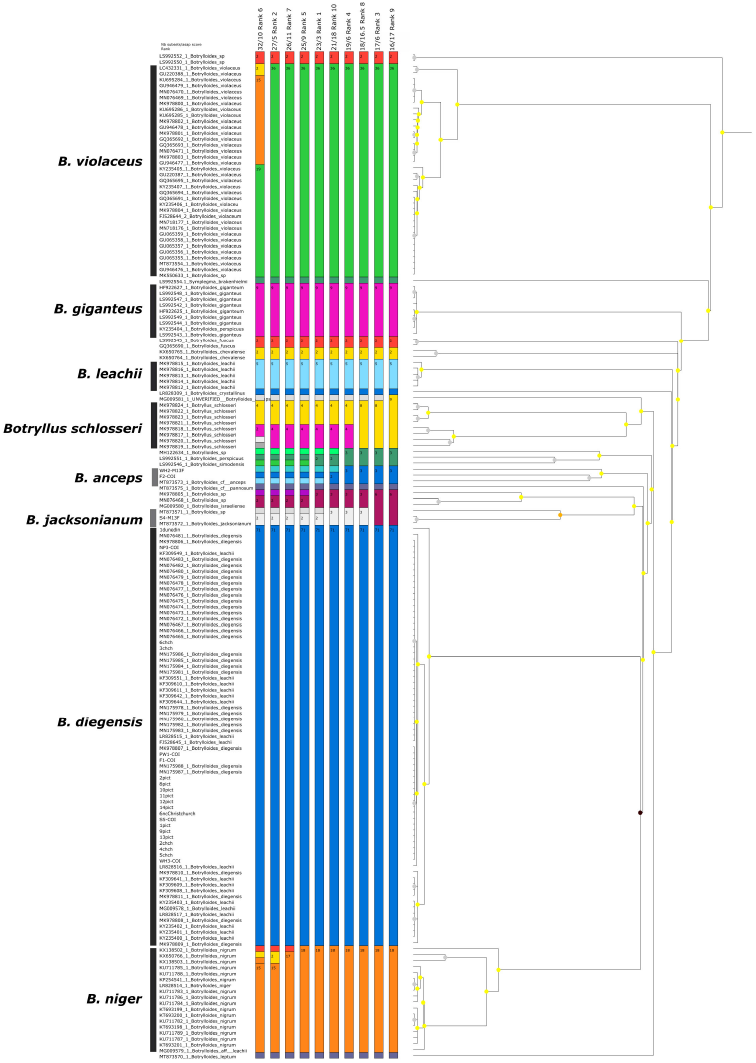
Molecular species delimitation using ASAP. ASAP scores were measured for *Botrylloides* species based on Jukes-cantor and kimura 2-parameter model selection. Colors illustrate the OTUs, while numbers within each OTU show the sample numbers belonging to that unit. Each column is a partition with the best ASAP score. The column headers are the OTU numbers and ASAP scores. The lowest ASAP score was taken into consideration for the OTU selection. The ultrametric clustering tree is shown on the right; node color indicates the p-value.

Seven haplotypes were found based on the clustering of the COI Network from the 7 sampling locations (Fig. 5). The distance values (0.2 - 1%) of the sequences support their identification as *B. diegensis* based on the COI species delineation threshold (< 2%) (Hebert et al., 2003) compared to database sequences based on BLAST mismatch ratio. Haplotype H1 appears to be widely distributed, found in both North and South Islands (Fig. 5). Additionally, three samples from Whangateau, Dunedin, and Invercargill were determined to be distinct species based on their evolutionary distances (Fig. 5; *B. aff. anceps* and *B. jacksonianum*), consistent with the phylogeny analysis (Fig. 3 and 4).

**Figure 5.**
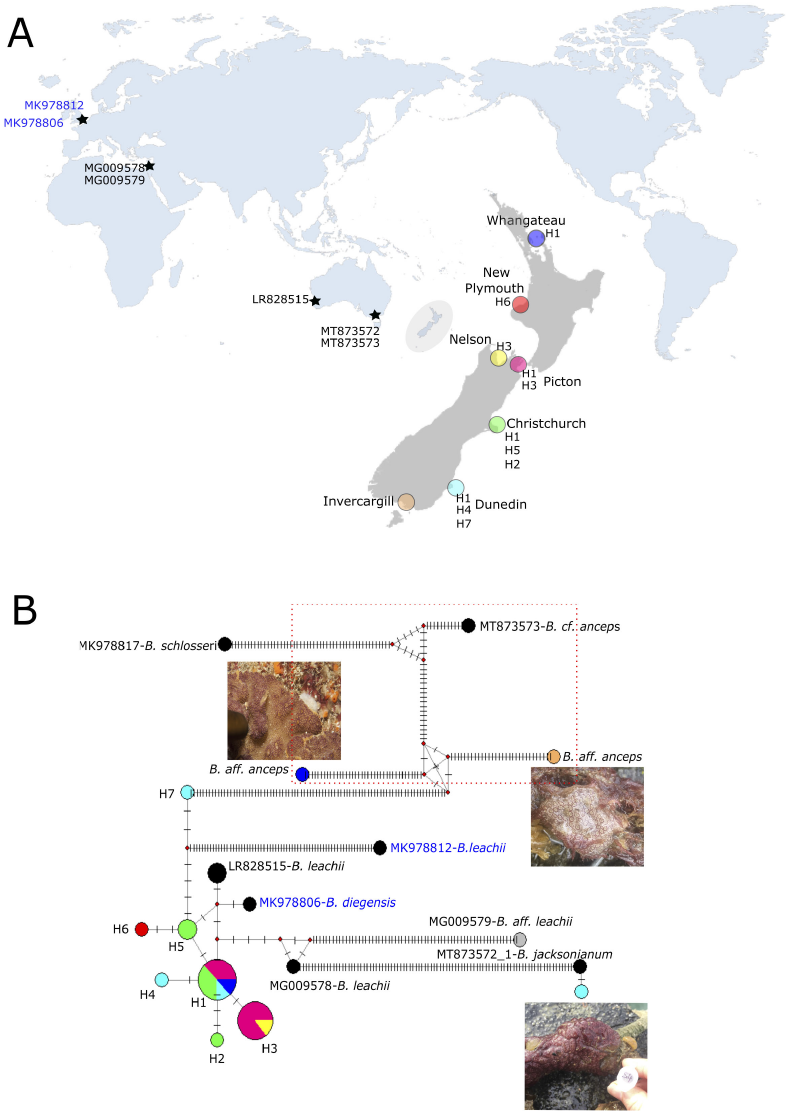
Phylogeography of the New Zealand Botryllid ascidians. (A) Location of sample sites in New Zealand (Turquoise: Dunedin, Green: Christchurch, Orange: Invercargill, Yellow: Nelson, Pink: Picton, Blue: Whangateau, Red: New Plymouth). The location of database sequences used in the analysis is also shown on the world map. (B) The strips on the lines indicate the mutation steps between the haplotypes. Circle size represents the frequency of the haplotype. Colors mark the regions where the samples were collected, as given on the map in panel A. Analyzed partial COI sequences are 647 bp. Grey and black circles indicate the database haplotypes. Black is for *B. diegensis/leachii*, grey is for *B. aff. leachii* (GenBank sequences). Blue text indicates *B. leachii* MK978812) and *B. diegensis* reference sequences (MK978806) (Viard et al., 2019). *B. anceps* cluster is highlighted with a red dashed-line box.

Besides mitochondrial marker COI, three other nuclear regions as H3, 18S, and 28S were analyzed. Based on H3, 18S, and 28S, Bayesian trees were constructed (Fig. 6). The H3 tree (Fig. 6A) confirms the clustering of the New Zealand samples with database sequence labelled as *B. leachii* (*B. diegensis* - Accession no: MG009592). The low evolutionary distance of database *B. leachii* sequences based on 18S (MG009583) and 28S (MG009588) barcodes further confirm the identification of current study sequences as *B. diegensis* (Fig. 6B, C). This *B. leachii* database sample is *B. diegensis* due to its COI barcode (MG009578) clustering with the *B. diegensis* sequences in the Bayesian phylogeny.

**Figure 6.**
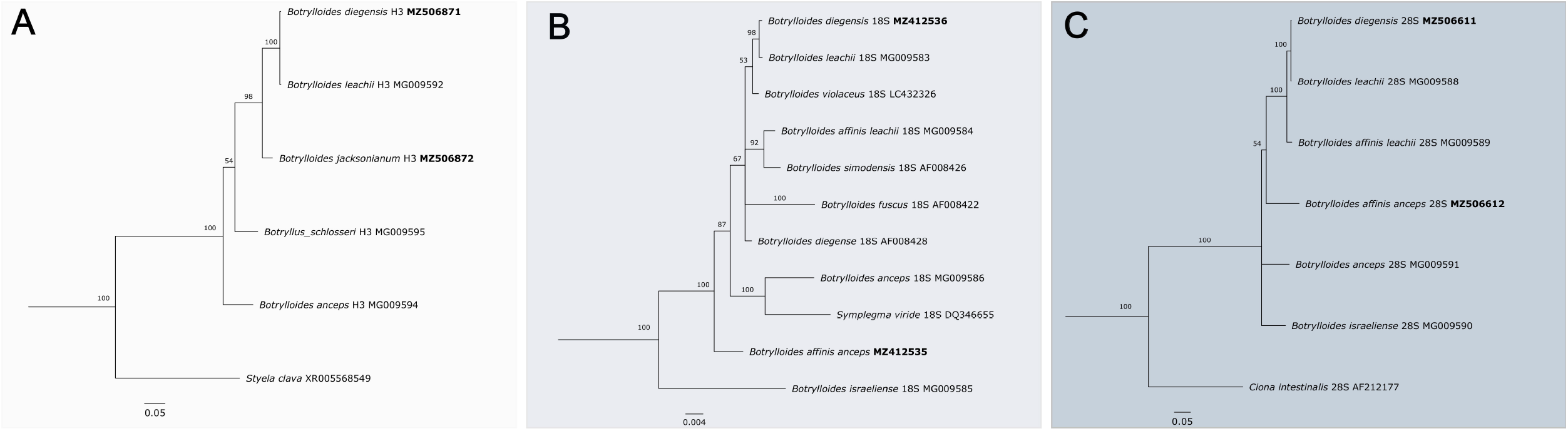
Bayesian trees based on nuclear markers. **A)** Tree is based on Histone 3 based on the Hasegawa-Kishino-Yano (HKY+G)) substitution model (Hasegawa et al., 1985). The outgroup is *Styela clava* (Accession no: XR_005568549). (B) Phylogeny based on 18S ribosomal subunit gene is constructed based on kimura 2-parameter with gamma (K80+G) substitution rate (Kimura, 1980). The outgroup is *Symplegma viride* (Accession no: DQ346655). **C)** 28S ribosomal subunit tree is calculated with general time reversible with gamma (GTR+G) model (Tavaré, 1986), *Ciona intestinalis* (AF212177) was used as the outgroup. Phylogenies are constructed with the study sequences and all database *Botrylloides* sequences from GenBank. Two independent runs were executed with Monte-Carlo Markov Chains. Ten million generations were measured, and the sampling frequency was 1000 for each generation. Split frequencies were lower than 0.01. Numbers on the tree indicate the bootstrap frequencies. The accession numbers of the sequences from this study are shown in bold.

Based on 18S, *B. aff. anceps* from Invercargill separated from the *Botrylloides anceps* from Israel (indicated as unverified on GenBank). The lack of 18S sequences in the database resulted in insufficient separation from the out-group (*Symplegma viride)* from the *Botrylloides* sequences. In contrast, the H3 and 28S out-group could separate in the tree further from the compared sequences. Phylogeny based on 28S assigned the *B. aff. anceps* from Whangateu more closely related to the *B. anceps* from Israel (Accession no: MG009591).

## DISCUSSION

*Botrylloides* colonies were collected from seven different regions of the North and South Island of New Zealand, and phylogenetic analysis was based on mitochondrial and nuclear markers. Due to the absence of the spoked wheel morphology and based on the comparative molecular results, most of the collected samples from New Zealand are identified as *Botrylloides diegensis* (Viard et al., 2019). For this reason, the previously reported studies from our lab that denoted our model organism as *Botrylloides leachii* are *Botrylloides diegensis* (Blanchoud, Rinkevich, et al., 2018; Blanchoud, Rutherford, et al., 2018; Blanchoud et al., 2017; Zondag, 2019; Zondag et al., 2016).

Two other *Botrylloides* species collected are the first molecular records from New Zealand: *Botrylloides jacksonianum* and *Botrylloides affinis anceps*. The species from Whangateau was assigned as *B. aff. anceps* due to its high molecular affinity to the record from Australia (MT873573) (Salonna et al., 2021). Although we could not obtain a detailed anatomical comparison for both these records, our result is consistent, particularly regarding *B. anceps*, as it is previously stated to be indigenous to the Indo-Pacific region (Brunetti, 2009). The presence of two larger lateral tentacles given for *B. anceps* from Israel also confirms the genetic affinity regarding our assignment (Brunetti, 2009). The ASAP analyses resulted in 3 separate OTUs for *B. anceps* samples (Fig. 4). Due to these samples’ high molecular and morphological affinity, we hypothesize *B. anceps* might be a species complex. Similarly, *Botyrllus schlosseri*, a species of the sister genus, was recently stated to be a species complex that contains divergent clades with high evolutionary distances (Ben-Hamo & Rinkevich, 2021; Brunetti et al., 2020; Brunetti et al., 2017; Nydam et al., 2017; Reem et al., 2021). A higher number of *B. anceps* samples would be needed to resolve this hypothesis.

Besides a single recent study (Salonna et al., 2021), little is known about *B. jacksonianum*. Salonna et al., (2021) concluded that *B. jacksonianum* (Herdman, 1886), which was synonymized with *B. leachii* by Kott (Kott, 1985, 2005), is a distinct species. Our network construction, phylogenetic tree and ABGD analyses also support the conclusions of this 2021 publication that it is a separate species. We found the zooid sizes of *B. diegensis* and *B. aff. anceps* is similar, while *B. jacksonianum* was larger. As a general observation, we commonly found *B. diegensis* during the summertime. However, *B. jacksonianum* was observed more often during the winter during our regular samplings in Dunedin.

As a single monophyletic cluster, we observed seven haplotypes from the New Zealand coasts based on COI sequences. Although a small number of the species was barcoded, a relatively higher number of haplotypic variations were recorded.

In conclusion, we could identify our samples using COI, H3,18S and 28S markers. As one of the first barcoding studies on *Botrylloides* ascidians from New Zealand, this study represents valuable insights into *Botrylloides* species diversity. We found three *Botrylloides* species - *Botrylloides diegensis, Botrylloides jacksonianum*, and *Botrylloides affinis anceps-* present in the marinas of Aotearoa New Zealand. For our previous publications, we checked our COI barcodes which matched at the time with the *B. leachii* sequences in the GenBank database and morphological information in the associated publications. However, since then, these sequences were determined as incorrectly annotated and belong to *B. diegensis*. Therefore, this study confirms that the species in our previous publications was not *B. leachii* but *B. diegensis*.

## ACKNOWLEDGEMENTS

We would like to thank Richard Taylor from the University of Auckland who kindly provided samples from Whangateau. We would also like to thank Dr. Michael Meier for the help during the samplings. Finally, we thank Stephanie Workman for her comments on earlier drafts. This study is supported by the Royal Society of New Zealand Marsden fund grant (UOO1713). B Temiz was supported by the Anatomy Department PhD scholarship from the University of Otago.

## DATA AVAILABILITY

The haplotype sequences are uploaded to the GenBank nucleotide database. The accession numbers are (MZ412536, MZ506611, MZ533117, MZ533119, MZ506871, MZ506872, MZ412535, MZ506612, MZ533118, MZ533120) (Table S2). The phylogenies are uploaded to the Open Tree of Life database (https://tree.opentreeoflife.org/curator/study/view/ot_2094). The accession numbers of the COI sequences and out-groups can be found in the Supplementary Information, Table S2 (current study sequences) and Table S3 (database sequences). The data generated during and/or analyzed during the current study are available from the corresponding author upon reasonable request.

## REFERENCES

Bandelt, H. J., Forster, P., & Rohl, A. (1999). Median-joining networks for inferring intraspecific phylogenies. Molecular Biology and Evolution, 16(1), 37–48. https://doi.org/10.1093/oxfordjournals.molbev.a026036

Belton, P. S., Tanner, S. F., Cartier, N., & Chanzy, H. (1989). High-Resolution Solid-State C-13 Nuclear Magnetic-Resonance Spectroscopy of Tunicin, an Animal Cellulose. Macromolecules, 22(4), 1615–1617. https://doi.org//10.1021/ma00194a019

Ben-Hamo, O., & Rinkevich, B. (2021). Botryllus schlosseri—A model colonial species in basic and applied studies. In Handbook of Marine Model Organisms in Experimental Biology (pp. 385–402). CRC Press.

Berrill, N. J. (1947). The developmental cycle of Botrylloides. Q J Microsc Sci, 88(Pt 4), 393–407. https://www.ncbi.nlm.nih.gov/pubmed/18904458

Blanchoud, S., Rinkevich, B., & Wilson, M. J. (2018). Whole-Body Regeneration in the Colonial Tunicate Botrylloides leachii. Results Probl Cell Differ, 65, 337–355. https://doi.org/10.1007/978-3-319-92486-1_16

Blanchoud, S., Rutherford, K., Zondag, L., Gemmell, N. J., & Wilson, M. J. (2018). De novo draft assembly of the Botrylloides leachii genome provides further insight into tunicate evolution. Sci Rep, 8(1), 5518. https://doi.org/10.1038/s41598-018-23749-w

Blanchoud, S., Zondag, L., Lamare, M. D., & Wilson, M. J. (2017). Hematological Analysis of the Ascidian Botrylloides leachii (Savigny, 1816) During Whole-Body Regeneration. Biol Bull, 232(3), 143–157. https://doi.org/10.1086/692841

Brunetti, R. (2009). Botryllid species (Tunicata, Ascidiacea) from the Mediterranean coast of Israel, with some considerations on the systematics of Botryllinae. Zootaxa (2289), 18–32. https://doi.org/10.11646/zootaxa.2289.1.2

Brunetti, R., Gissi, C., Pennati, R., Caicci, F., Gasparini, F., & Manni, L. (2015). Morphological evidence that the molecularly determined Ciona intestinalis type A and type B are different species: Ciona robusta and Ciona intestinalis. Journal of Zoological Systematics and Evolutionary Research, 53(3), 186–193. https://doi.org/10.1111/jzs.12101

Brunetti, R., Griggio, F., Mastrototaro, F., Gasparini, F., & Gissi, C. (2020). Toward a resolution of the cosmopolitan Botryllus schlosseri species complex (Ascidiacea, Styelidae): mitogenomics and morphology of clade E (Botryllus gaiae). Zoological Journal of the Linnean Society, 190(4), 1175–1192.

Brunetti, R., Manni, L., Mastrototaro, F., Gissi, C., & Gasparini, F. (2017). Fixation, description and DNA barcode of a neotype for Botryllus schlosseri Pallas, 1766)(Tunicata, Ascidiacea). Zootaxa, 4353(1), 29–50. https://doi.org/10.11646/zootaxa.4353.1.2

Brunetti, R., & Mastrototaro, F. (2012). Botrylloides pizoni, a new species of Botryllinae (Ascidiacea) from the Mediterranean Sea. Zootaxa, 36, 28–36. https://www.marinespecies.org/aphia.php?p=sourcedetails&id=164338

Burighel, P., Brunetti, R., & Zaniolo, G. (1976). Hibernation of the Colonial Ascidian Botrylloides Leachi (Savigny): Histological Observations. Bollettino di zoologia, 43(3), 293–301. https://doi.org/10.1080/11250007609430146

Carlton, J. T. (2009). Deep invasion ecology and the assembly of communities in historical time. Springer. https://doi.org/10.1007/978-3-540-79236-9_2

Carman, M. R., Morris, J. A., Karney, R. C., & Grunden, D. W. (2010). An initial assessment of native and invasive tunicates in shellfish aquaculture of the North American east coast. Journal of Applied Ichthyology, 26, 8–11. https://doi.org/10.1111/j.1439-0426.2010.01495.x

Darriba, D., Taboada, G. L., Doallo, R., & Posada, D. (2012). jModelTest 2: more models, new heuristics and parallel computing. Nature methods, 9(8), 772–772.

Delsuc, F., Brinkmann, H., Chourrout, D., & Philippe, H. (2006). Tunicates and not cephalochordates are the closest living relatives of vertebrates. Nature, 439(7079), 965–968. https://doi.org/10.1038/nature04336

Folmer, O., Black, M., Hoeh, W., Lutz, R., & Vrijenhoek, R. (1994). DNA primers for amplification of mitochondrial cytochrome c oxidase subunit I from diverse metazoan invertebrates. Mol Mar Biol Biotechnol, 3(5), 294–299.

Gemmell, N. J., & Akiyama, S. (1996). An efficient method for the extraction of DNA from vertebrate tissues. Trends Genet, 12(9), 338–339. https://doi.org/10.1016/s0168-9525(96)80005-9

Hall, T. A. (1999). BioEdit: A User-Friendly Biological Sequence Alignment Editor and Analysis Program for Windows

Hasegawa, M., Kishino, H., & Yano, T.-a. (1985). Dating of the human-ape splitting by a molecular clock of mitochondrial DNA. Journal of molecular evolution, 22(2), 160–174.

Hebert, P. D., Ratnasingham, S., & deWaard, J. R. (2003). Barcoding animal life: cytochrome c oxidase subunit 1 divergences among closely related species. Proc Biol Sci, 270 Suppl 1, S96–99. https://doi.org/10.1098/rsbl.2003.0025

Herdman, W. A. (1886). Report on the Tunicata collected during the years 1873-1876. Part 2, Ascidiae compositae. Zool. Chall. Exp., 14(38), 1–425. https://www.marinespecies.org/aphia.php?p=sourcedetails&id=43289

Herdman, W. A. (1891). A Revised Classification of the Tunicata, with Definitions of the Orders, Suborders, Families, Subfamilies, and Genera, and Analytical Keys to the Species. Journal of the Linnean Society of London, Zoology, 23(148), 558–652. https://doi.org/10.1111/j.1096-3642.1891.tb02403.x

Herdman, W. A. (1899). Descriptive catalogue of the tunicata in the Australian Museum Sydney N.S.W. Austr. Mus. Sydney Catal., 17, 1–139. https://www.marinespecies.org/aphia.php?p=sourcedetails&id=43294

Jukes, T. H., & Cantor, C. R. (1969). Evolution of protein molecules. Mammalian protein metabolism, 3, 21–132.

Karahan, A., Douek, J., Paz, G., & Rinkevich, B. (2016). Population genetics features for persistent, but transient, Botryllus schlosseri (Urochordata) congregations in a central Californian marina. Molecular Phylogenetics and Evolution, 101, 19–31. https://doi.org/10.1016/j.ympev.2016.05.005

Karahan, A., Öztürk, E., Temiz, B., & Blanchoud, S. (2022). Studying TunicataTunicata WBRWhole-body regeneration (WBR)Using Botrylloides anceps. In S. Blanchoud & B. Galliot (Eds.), Whole-Body Regeneration: Methods and Protocols (pp. 311–332). Springer US. https://doi.org/10.1007/978-1-0716-2172-1_16

Kimura, A. (1980). Statistical properties of random wave groups. Coastal Engineering Proceedings (17), 175–175.

Kocot, K. M., Tassia, M. G., Halanych, K. M., & Swalla, B. J. (2018). Phylogenomics offers resolution of major tunicate relationships. Molecular Phylogenetics and Evolution, 121, 166–173. https://doi.org/10.1016/j.ympev.2018.01.005

Kott, P. (1985). The Australian Ascidiacea. Part 1, Phlebobranchia and Stolidobranchi. In Memoirs of the Queensland Museum (Vol. 23, pp. 1–439). Queensland Museum,.

Kott, P. (2005). Catalogue of Tunicata in Australian waters/P. Kott. Australian Biological Resources Study.

Manni, L., Anselmi, C., Cima, F., Gasparini, F., Voskoboynik, A., Martini, M., Peronato, A., Burighel, P., Zaniolo, G., & Ballarin, L. (2019). Sixty years of experimental studies on the blastogenesis of the colonial tunicate Botryllus schlosseri. Developmental Biology, 448(2), 293–308. https://doi.org/10.1016/j.ydbio.2018.09.009

Miralles, L., Ardura, A., Arias, A., Borrell, Y. J., Clusa, L., Dopico, E., de Rojas, A. H., Lopez, B., Munoz-Colmenero, M., Roca, A., Valiente, A. G., Zaiko, A., & Garcia-Vazquez, E. (2016). Barcodes of marine invertebrates from north Iberian ports: Native diversity and resistance to biological invasions. Marine Pollution Bulletin, 112(1-2), 183–188. https://doi.org/10.1016/j.marpolbul.2016.08.022

Nydam, M. L., Giesbrecht, K. B., & Stephenson, E. E. (2017). Origin and dispersal history of two colonial ascidian clades in the Botryllus schlosseri species complex. PLoS One, 12(1), e0169944.

Oka, A. (1927). Zur kenntnis der japanishen Botryllidae (Vorlaufige Mitteilung). Proc. Imp. Acad, 3, 607–609. https://www.marinespecies.org/aphia.php?p=sourcedetails&id=43591

Oka, A. (1928). Uber eine merkwurdige Botryllus-Art, B. primigenus nov. sp. Proc. Imp. Acad. Tokyo, 6, 303–305. https://www.marinespecies.org/aphia.php?p=sourcedetails&id=43592

Page, M., & Kelly, M. (2013). Awesome Ascidians: A Guide to the Sea Squirts of New Zealand. NIWA, Wellington.

Page, M. J., Willis, T. J., & Handley, S. J. (2014). The colonial ascidian fauna of Fiordland, New Zealand, with a description of two new species. Journal of Natural History, 48(27-28), 1653–1688. https://doi.org/10.1080/00222933.2014.896487

Pallas, P. S. (1766). Elenchus zoophytorum sistens generum adumbrationes generaliores et specierum cognitarum succintas descriptiones, cum selectis auctorum synonymis. [book]. https://doi.org/10.5962/bhl.title.6595

Pérès, J. M. (1949). Contribution à l’étude des Ascidies de la côte occidentale d’Afrique. Bulletin de l’Institut français d’Afrique noire, 11, 159–207. https://www.marinespecies.org/aphia.php?p=sourcedetails&id=43625

Puillandre, N., Brouillet, S., & Achaz, G. (2021). ASAP: assemble species by automatic partitioning. Molecular Ecology Resources, 21(2), 609–620.

Rambaut, A. (2018). FigTree v1.4.4. https://github.com/rambaut/figtree.

Reem, E., Douek, J., & Rinkevich, B. (2018). Ambiguities in the taxonomic assignment and species delineation of botryllid ascidians from the Israeli Mediterranean and other coastlines. Mitochondrial DNA Part A, 29(7), 1073–1080. https://doi.org/10.1080/24701394.2017.1404047

Reem, E., Douek, J., & Rinkevich, B. (2021). A critical deliberation of the ‘species complex’status of the globally spread colonial ascidian Botryllus schlosseri. Journal of the Marine Biological Association of the United Kingdom, 101(7), 1047–1060.

Rinkevich, B. (2005). Natural chimerism in colonial urochordates. Journal of Experimental Marine Biology and Ecology, 322(2), 93–109. https://doi.org/10.1016/j.jembe.2005.02.020

Rinkevich, B., Shlemberg, Z., & Fishelson, L. (1995). Whole-body protochordate regeneration from totipotent blood cells. Proc Natl Acad Sci U S A, 92(17), 7695–7699. https://doi.org/10.1073/pnas.92.17.7695

Rinkevich, Y., Douek, J., Haber, O., Rinkevich, B., & Reshef, R. (2007). Urochordate whole body regeneration inaugurates a diverse innate immune signaling profile. Developmental Biology, 312(1), 131–146. https://doi.org/10.1016/j.ydbio.2007.09.005

Ritter, W. E., & Forsyth, R. H. (1917). Ascidian of the littoral zone of southern California. Univ. California Publ., Zool., 16, 439–512. https://www.marinespecies.org/aphia.php?p=sourcedetails&id=43674

Rocha, R. M., Salonna, M., Griggio, F., Ekins, M., Lambert, G., Mastrototaro, F., Fidler, A., & Gissi, C. (2019). The power of combined molecular and morphological analyses for the genus Botrylloides: identification of a potentially global invasive ascidian and description of a new species. Systematics and Biodiversity, 17(5), 509–526. https://doi.org/10.1080/14772000.2019.1649738

Ronquist, F., Teslenko, M., van der Mark, P., Ayres, D. L., Darling, A., Hohna, S., Larget, B., Liu, L., Suchard, M. A., & Huelsenbeck, J. P. (2012). MrBayes 3.2: efficient Bayesian phylogenetic inference and model choice across a large model space. Syst Biol, 61(3), 539–542. https://doi.org/10.1093/sysbio/sys029

Rosner, A., Kravchenko, O., & Rinkevich, B. (2019). IAP genes partake weighty roles in the astogeny and whole body regeneration in the colonial urochordate Botryllus schlosseri. Developmental Biology, 448(2), 320–341. https://doi.org/10.1016/j.ydbio.2018.10.015

Salonna, M., Gasparini, F., Huchon, D., Montesanto, F., Haddas-Sasson, M., Ekins, M., McNamara, M., Mastrototaro, F., & Gissi, C. (2021). An elongated COI fragment to discriminate botryllid species and as an improved ascidian DNA barcode. Sci Rep, 11(1), 4078. https://doi.org/10.1038/s41598-021-83127-x

Shenkar, N., & Swalla, B. J. (2011). Global diversity of Ascidiacea. PLoS One, 6(6), e20657. https://doi.org/10.1371/journal.pone.0020657

Stefaniak, L., Lambert, G., Gittenberger, A., Zhang, H., Lin, S., & Whitlatch, R. B. (2009). Genetic conspecificity of the worldwide populations of Didemnum vexillum Kott, 2002. Aquatic Invasions, 4(1), 29–44.

Tavaré, S. (1986). Some probabilistic and statistical problems in the analysis of DNA sequences. Lectures on mathematics in the life sciences, 17(2), 57–86.

Thompson, J. D., Gibson, T. J., Plewniak, F., Jeanmougin, F., & Higgins, D. G. (1997). The CLUSTAL_X windows interface: flexible strategies for multiple sequence alignment aided by quality analysis tools. Nucleic acids research, 25(24), 4876–4882.

Viard, F., Roby, C., Turon, X., Bouchemousse, S., & Bishop, J. (2019). Cryptic Diversity and Database Errors Challenge Non-indigenous Species Surveys: An Illustration With Botrylloides spp. in the English Channel and Mediterranean Sea [Original Research]. Frontiers in Marine Science, 6. https://doi.org/10.3389/fmars.2019.00615

Zhan, A., Briski, E., Bock, D. G., Ghabooli, S., & MacIsaac, H. J. (2015). Ascidians as models for studying invasion success. Marine Biology, 162(12), 2449–2470. https://doi.org/10.1007/s00227-015-2734-5

Zondag, L., Clarke, R., Wilson, M. J. (2019). Histone deacetylase activity is required for Botrylloides leachii whole-body regeneration. J Exp Biol, 222(Pt 15). https://doi.org/10.1242/jeb.203620

Zondag, L., Rutherford, K., Gemmell, N. J., & Wilson, M. J. (2016). Uncovering the pathways underlying whole body regeneration in a chordate model, Botrylloides leachi using de novo transcriptome analysis. BMC Genomics, 17, 114. https://doi.org/10.1186/s12864-016-2435-6

