## Supplementary Information for "Identification and characterization of *Botrylloides* (Styelidae) species from Aotearoa New Zealand coasts"

| | Accession Number | Species |
|---|---|---|
| 1 | FJ528644.2 | *Botrylloides violaceum* |
| 2 | FJ528645.1 | *Botrylloides leachi* |
| 3 | GQ365690.1 | *Botrylloides fuscus* |
| 4 | GQ365691.1 | *Botrylloides violaceus* |
| 5 | GQ365692.1 | *Botrylloides violaceus* |
| 6 | GQ365693.1 | *Botrylloides violaceus* |
| 7 | GQ365694.1 | *Botrylloides violaceus* |
| 8 | GQ365695.1 | *Botrylloides violaceus* |
| 9 | GU065355.1 | *Botrylloides violaceus* |
| 10 | GU065356.1 | *Botrylloides violaceus* |
| 11 | GU065357.1 | *Botrylloides violaceus* |
| 12 | GU065358.1 | *Botrylloides violaceus* |
| 13 | GU065359.1 | *Botrylloides violaceus* |
| 14 | GU220387.1 | *Botrylloides violaceus* |
| 15 | GU220388.1 | *Botrylloides violaceus* |
| 16 | GU946476.1 | *Botrylloides violaceus* |
| 17 | GU946477.1 | *Botrylloides violaceus* |
| 18 | GU946478.1 | *Botrylloides violaceus* |
| 19 | GU946479.1 | *Botrylloides violaceus* |
| 20 | HF922625.1 | *Botrylloides giganteum* |
| 21 | HF922626.2 | *Botrylloides giganteum* |
| 22 | HF922627.1 | *Botrylloides giganteum* |
| 23 | KF309549.1 | *Botrylloides leachii* |
| 24 | KF309551.1 | *Botrylloides leachii* |
| 25 | KF309608.1 | *Botrylloides leachii* |
| 26 | KF309609.1 | *Botrylloides leachii* |
| 27 | KF309610.1 | *Botrylloides leachii* |
| 28 | KF309611.1 | *Botrylloides leachii* |
| 29 | KF309641.1 | *Botrylloides leachii* |
| 30 | KF309642.1 | *Botrylloides leachii* |
| 31 | KF309644.1 | *Botrylloides leachii* |
| 32 | KP254541.1 | *Botrylloides nigrum* |
| 33 | KT693198.1 | *Botrylloides nigrum* |
| 34 | KT693199.1 | *Botrylloides nigrum* |
| 35 | KT693200.1 | *Botrylloides nigrum* |
| 36 | KT693201.1 | *Botrylloides nigrum* |
| 37 | KU695284.1 | *Botrylloides violaceus* |
| 38 | KU695285.1 | *Botrylloides violaceus* |
| 39 | KU695286.1 | *Botrylloides violaceus* |
| 40 | KU711782.1 | *Botrylloides nigrum* |
| 41 | KU711783.1 | *Botrylloides nigrum* |
| 42 | KU711784.1 | *Botrylloides nigrum* |
| 43 | KU711785.1 | *Botrylloides nigrum* |
| 44 | KU711786.1 | *Botrylloides nigrum* |
| 45 | KU711787.1 | *Botrylloides nigrum* |
| 46 | KU711788.1 | *Botrylloides nigrum* |
| 47 | KU711789.1 | *Botrylloides nigrum* |

| | | |
|---|---|---|
| **48** | KX138502.1 | *Botrylloides nigrum* |
| **49** | KX138503.1 | *Botrylloides nigrum* |
| **50** | KX650764.1 | *Botrylloides chevalense* |
| **51** | KX650765.1 | *Botrylloides chevalense* |
| **52** | KX650766.1 | *Botrylloides nigrum* |
| **53** | KY235400.1 | *Botrylloides leachii* |
| **54** | KY235401.1 | *Botrylloides leachii* |
| **55** | KY235402.1 | *Botrylloides leachii* |
| **56** | KY235403.1 | *Botrylloides leachii* |
| **57** | KY235404.1 | *Botrylloides perspicuus* |
| **58** | KY235405.1 | *Botrylloides violaceus* |
| **59** | KY235406.1 | *Botrylloides violaceus* |
| **60** | KY235407.1 | *Botrylloides violaceus* |
| **61** | LC432331.1 | *Botrylloides violaceus* |
| **62** | LR828309.1 | *Botrylloides crystallinus* |
| **63** | LR828514.1 | *Botrylloides niger* |
| **64** | LR828515.1 | *Botrylloides leachii* |
| **65** | LR828516.1 | *Botrylloides leachii* |
| **66** | LR828517.1 | *Botrylloides leachii* |
| **67** | LS992542.1 | *Botrylloides giganteus* |
| **68** | LS992543.1 | *Botrylloides giganteus* |
| **69** | LS992544.1 | *Botrylloides giganteus* |
| **70** | LS992545.1 | *Botrylloides fuscus* |
| **71** | LS992546.1 | *Botrylloides simodensis* |
| **72** | LS992547.1 | *Botrylloides giganteus* |
| **73** | LS992548.1 | *Botrylloides giganteus* |
| **74** | LS992549.1 | *Botrylloides giganteus* |
| **75** | LS992550.1 | *Botrylloides sp.* |
| **76** | LS992551.1 | *Botrylloides perspicuus* |
| **77** | LS992552.1 | *Botrylloides sp.* |
| **78** | MG009578.1 | *Botrylloides leachii* |
| **79** | MG009579.1 | *Botrylloides aff. leachii* |
| **80** | MG009580.1 | *Botrylloides israeliense* |
| **81** | MG009581.1 | *Botrylloides anceps* |
| **82** | MH122634.1 | *Botrylloides sp.* |
| **83** | MK550633.1 | *Botrylloides sp.* |
| **84** | MK978800.1 | *Botrylloides violaceus* |
| **85** | MK978801.1 | *Botrylloides violaceus* |
| **86** | MK978802.1 | *Botrylloides violaceus* |
| **87** | MK978803.1 | *Botrylloides violaceus* |
| **88** | MK978804.1 | *Botrylloides violaceus* |
| **89** | MK978805.1 | *Botrylloides sp.* |
| **90** | MK978806.1 | *Botrylloides diegensis* |
| **91** | MK978807.1 | *Botrylloides diegensis* |
| **92** | MK978808.1 | *Botrylloides diegensis* |
| **93** | MK978809.1 | *Botrylloides diegensis* |
| **94** | MK978810.1 | *Botrylloides diegensis* |
| **95** | MK978811.1 | *Botrylloides diegensis* |

| | | |
|---|---|---|
| **96** | MK978812.1 | *Botrylloides leachii* |
| **97** | MK978813.1 | *Botrylloides leachii* |
| **98** | MK978814.1 | *Botrylloides leachii* |
| **99** | MK978815.1 | *Botrylloides leachii* |
| **100** | MK978816.1 | *Botrylloides leachii* |
| **101** | MK978817.1 | *Botryllus schlosseri* |
| **102** | MK978818.1 | *Botryllus schlosseri* |
| **103** | MK978819.1 | *Botryllus schlosseri* |
| **104** | MK978820.1 | *Botryllus schlosseri* |
| **105** | MK978821.1 | *Botryllus schlosseri* |
| **106** | MK978822.1 | *Botryllus schlosseri* |
| **107** | MK978823.1 | *Botryllus schlosseri* |
| **108** | MK978824.1 | *Botryllus schlosseri* |
| **109** | MN076465.1 | *Botrylloides diegensis* |
| **110** | MN076466.1 | *Botrylloides diegensis* |
| **111** | MN076467.1 | *Botrylloides diegensis* |
| **112** | MN076468.1 | *Botrylloides sp.* |
| **113** | MN076469.1 | *Botrylloides violaceus* |
| **114** | MN076470.1 | *Botrylloides violaceus* |
| **115** | MN076471.1 | *Botrylloides violaceus* |
| **116** | MN076472.1 | *Botrylloides diegensis* |
| **117** | MN076473.1 | *Botrylloides diegensis* |
| **118** | MN076474.1 | *Botrylloides diegensis* |
| **119** | MN076475.1 | *Botrylloides diegensis* |
| **120** | MN076476.1 | *Botrylloides diegensis* |
| **121** | MN076477.1 | *Botrylloides diegensis* |
| **122** | MN076478.1 | *Botrylloides diegensis* |
| **123** | MN076479.1 | *Botrylloides diegensis* |
| **124** | MN076480.1 | *Botrylloides diegensis* |
| **125** | MN076481.1 | *Botrylloides diegensis* |
| **126** | MN076482.1 | *Botrylloides diegensis* |
| **127** | MN076483.1 | *Botrylloides diegensis* |
| **128** | MN175978.1 | *Botrylloides diegensis* |
| **129** | MN175979.1 | *Botrylloides diegensis* |
| **130** | MN175980.1 | *Botrylloides diegensis* |
| **131** | MN175981.1 | *Botrylloides diegensis* |
| **132** | MN175982.1 | *Botrylloides diegensis* |
| **133** | MN175983.1 | *Botrylloides diegensis* |
| **134** | MN175984.1 | *Botrylloides diegensis* |
| **135** | MN175985.1 | *Botrylloides diegensis* |
| **136** | MN175986.1 | *Botrylloides diegensis* |
| **137** | MN175987.1 | *Botrylloides diegensis* |
| **138** | MN175988.1 | *Botrylloides diegensis* |
| **139** | MN718176.1 | *Botrylloides violaceus* |
| **140** | MN718177.1 | *Botrylloides violaceus* |
| **141** | MT637977.1 | *Botrylloides sp.* |
| **142** | MT738742.1 | *Botrylloides simodensis* |
| **143** | MT738743.1 | *Botrylloides simodensis* |

| | | |
|---|---|---|
| **144** | MT873554.1 | *Botrylloides violaceus* |
| **145** | MT873570.1 | *Botrylloides leptum* |
| **146** | MT873571.1 | *Botrylloides sp.* |
| **147** | MT873572.1 | *Botrylloides jacksonianum* |
| **148** | MT873573.1 | *Botrylloides cf. anceps* |
| **149** | MT873575.1 | *Botrylloides cf. pannosum* |
